# The complete genome sequence of *Pseudomonas syringae* pv. *actinidifoliorum* ICMP 18803

**DOI:** 10.1101/2022.10.03.510724

**Authors:** Matthew D. Templeton, Saadiah Arshed, Mark T. Andersen, Jay Jayaraman

## Abstract

The complete genome of *Pseudomonas syringae* pv. *actinidifoliorum* ICMP18803 (Pfm) was sequenced using the Oxford Nanopore minION platform to an average read depth of 123. The genome assembled into a single chromosome of 6,353,853 bp after error-correction with Illumina short reads using Pilon. The complement of effector genes from a *P. syringae* pathovar plays the predominant role in defining its pathogenicity. Automatic gene annotation pipelines often poorly identify and name effector genes, however. Despite Pfm being a relatively weak pathogen of kiwifruit, a set of 31 effectors, 26 of which were full length, was identified by mapping the comprehensive effector library generated by Dillon et al. (2019). The Pfm genome with the effector complement, correctly named and annotated was resubmitted to Genbank (CP081457).

---

*Pseudomonas syringae* pv. *actinidifoliorum* (Pfm) was first isolated and characterised from kiwifruit orchards in New Zealand in 2010 (Chapman et al. 2012). It was initially classified as *Pseudomonas syringae* pv. *actinidiae* (Psa) group 4 (or LV for low virulence) due to the high degree of similarity between sequences used for multi-locus sequence analysis (MLSA) between it and other Psa strains (Chapman et al. 2012). Extensive pathogenicity testing of Pfm isolates using several different assay methods determined that this collection of strains caused a distinct and milder set of symptoms on kiwifruit compared to Psa. In contrast to the highly aggressive isolates of Psa biovar 3 (Psa3) which cause leaf-spotting, cane dieback and weeping trunk cankers, particularly on cultivars of *Actinidia chinensis* var *chinensis*, Pfm symptomology is largely restricted to mild leaf spotting (Ferrante and Scortichini 2015; Vanneste et al. 2013). Furthermore, *in planta* bacterial growth and movement assays revealed that Pfm grew to a maximum of 10^5^-10^6^ colony forming units per square centimetre, as opposed to 10^8^ for Psa3 and was capable of only limited systemic movement (Jayaraman et al. 2020; McAtee et al. 2018; McCann et al. 2013). For these reasons, Pfm was given its own pathovar designation (Cunty et al. 2015).

Pfm has been found to have a widespread distribution throughout kiwifruit growing regions in Europe, Australasia and Asia (Abelleira et al. 2015; Cunty et al. 2015; Vanneste et al. 2013). A phylogenetic tree generated from a multi-locus sequence analysis revealed significant genetic variation between these isolates which resolved into four lineages (Cunty et al. 2015). This suggests that there has been a long-term association between Pfm and *Actinidia* spp. that precedes the international outbreak of its more pathogenic relative Psa biovar 3.

Several groups have released short-read assemblies of various Pfm isolates (Butler et al. 2013; Cunty et al. 2016; McCann et al. 2013). In this paper we report the complete sequence of Pfm ICMP18803 isolated from Hawke’s Bay, New Zealand in 2010 (PRJNA167409; SAMN13855180). Genomic DNA from Pfm ICMP18803 was purified using a GenePure kit from Qiagen (Hilden, Germany) as described in McCann et al. (2013). Purified DNA was sequenced using the Oxford Nanopore minION platform to a read depth of 123 fold. The genome was assembled into a single contig using Flye 2.7.1, and error-corrected with Illumina short reads using Pilon 1.23 (Kolmogorov et al. 2020; Walker et al. 2014). All suggested short indel corrections were accepted. The resulting single chromosome was 6,353,853 bp in length. The genome was annotated using the PGAAP pipeline (Li et al. 2021). Nanopore and Illumina reads were deposited to the Sequence Read Archive (PRJNA167409).

Although isolates of Pfm presented only weak symptoms on kiwifruit, the genome has 31 effectors, of which 26 are full-length and expected to be active, plus secondary metabolite pathways which might contribute to pathogenicity (Tables 1 and 2). The PGAAP and other automatic annotation pipelines often annotate *P. syringae* effectors incorrectly. The Pfm effector complement was manually curated and annotated using the nomenclature used by Dillon et al. (2019). This was achieved by mapping all effectors from *P. syringae* in supplementary file 3 from Dillon et al. (2019) to the Pfm 18803 genome using the map to reference function in Geneious 10 (Biomatters). Effector alleles with 100% homology to the Pfm ICMP18803 genome were used to accurately annotate the start and stop sites and assign the correct name of the effector family (Table 1). One previously undiscovered effector that formed a new clade in the AvrPto1 phylogeny, AvrPto1r, was identified. This effector was co-located with HopF1b, HopAR1e, HopAF1b and HopAB1e which may be equivalent to the Exchangeable Effector Locus (EEL) in *P. syringae* pv. *tomato* and Psa, although the EEL is in a different location from the Conserved Effector Locus and the Type Three Secretion System (Alfano et al. 2000; McCann et al. 2013).

**Table 1.** The effector complement of *Pseudomonas syringae* pv. *actinidifoliorum* ICMP18803. Effectors were identified using the comprehensive *P. syringae* effector library compiled by Dillon et al. (2019). These genes were mapped to the *Pseudomonas syringae* pv. *actinidifoliorum* genome in Geneious (https://www.geneious.com) using the map to reference function. This was used to determine the correct stop and start sites and to assign the correct family designation for each effector.

| Effector | locus tag | Old name | Full CDS | Chaperone | Hrp boxL |
| --- | --- | --- | --- | --- | --- |
| HopY1d | A237_00695 | HopY1 | premature stop | n/a | yes |
| HopBO1c | A237_00840 | HopX2 | yes | n/a | yes |
| HopAS1b | A237_02165 | HopAS1 | yes | n/a | yes |
| HopW1f | A237_04025 | HopW1 | yes | n/a | yes |
| HopR1b | A237_04035 | HopR1 | yes | n/a | yes |
| HopAG1d | A237_04175 | HopAG1 | yes | A237_04173 | yes |
| HopAH1k | A237_04180 | HopAH1 | yes | n/a | yes (HopAG1) |
| HopAI1e | A237_04185 | HopAI1 | yes | n/a | yes (HopAG1) |
| HopF1b | A237_05265 | HopF1 | yes | A237_05260 | yes |
| HopAR1e | A237_05275 | HopAR1 | yes | n/a | yes |
| AvrPto1r | A237_05285 | N/A | premature stop | n/a | yes |
| HopAF1b | A237_05320 | HopAF1 | yes | n/a | yes (HopAB1) |
| HopAB1e | A237_05325 | HopAY1 | yes | n/a | yes |
| HopN1a | A237_06825 | HopN1 | yes | A237_06820 | yes |
| HopAA1d | A237_06835 | HopAA1 | yes | n/a | yes |
| HopM1f | A237_06850 | HopM1 | yes | A237_06845 | yes |
| AvrE1d | A237_06865 | AvrE1 | yes | A237_06855 | yes |
| HopX1d | A237_07010 | HopX1 | yes | n/a | yes (HrpK1) |
| HopAZ1a | A237_09620 | HopAZ1 | yes | n/a | yes |
| HopAH1a | A237_11260 | HopAH2-1 | yes | n/a | no |
| HopAH1i | A237_11265-75 | HopAH2-2 | Tn inactivated | n/a | no |
| HopW1c | A237_11645 | HopW1 | yes | n/a | yes |
| HopAB1i | A237_12225 | HopAB3 | yes | n/a | yes |
| HopE1a | A237_18035 | HopE1 | yes | n/a | yes |
| HopAF1f | A237_20845 | HopAF1-2 | yes | n/a | yes |
| HopS2c | A237_23320 | HopS2 | yes | A237_23325 | yes |
| HopT1c | A237_23330 | HopT1 | yes | n/a | yes (HopO2b) |
| HopO1a | A237_23335 | HopO1 | yes | n/a | yes (HopO2b) |
| HopO2b | A237_23340-50 | HopS1 | Tn inactivated | A237_23355 | yes |
| HopI1c | A237_23700 | HopI1 | premature stop | n/a | yes |
| HopA1a | A237_26495 | HopA1 | yes | A237_26490 | yes |

**Table 2.**
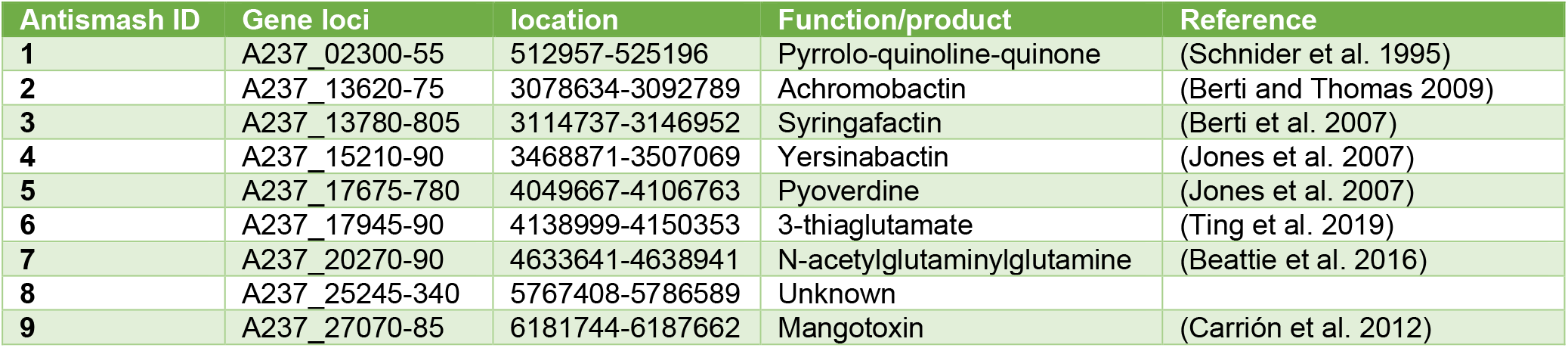
List of biosynthetic pathways identified using AntiSMASH.

The effector complement from different isolates or lineages within a *P. syringae* pathovar often varies, for example within biovars of Psa (McCann et al. 2013). This was also the case for Pfm (Figure 1). Pfm isolates from Europe and New Zealand largely had a conserved effector complement. The more recently discovered isolates from Japan, which form two new lineages, however were missing up to seven of these effectors (Figure 1). Examination of the regions around these effectors suggested that they were lost or acquired via elements associated with horizontal gene transfer.

**Figure 1.**
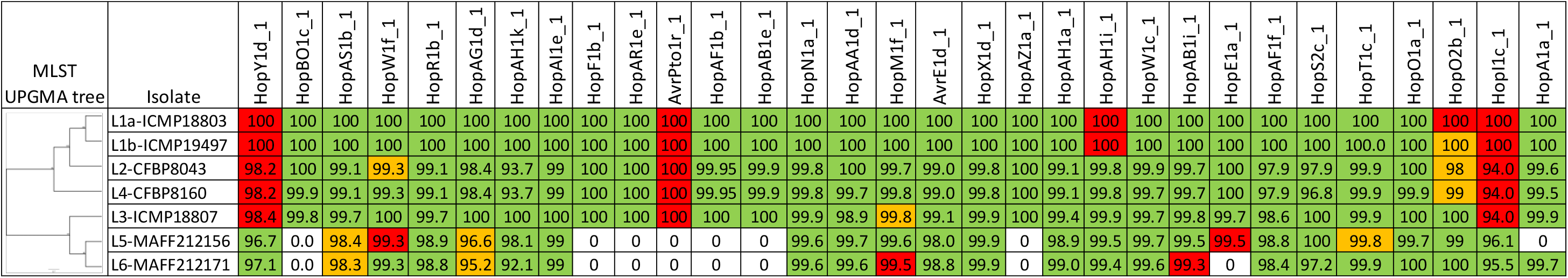
Comparison of the effector complement from different lineages of *Pseudomonas syringae* pv. *actinidifoliorum*. BLAST was used to search representative genomes from each lineage of *Pseudomonas syringae* pv. *actinidifoliorum* (Pfm) with every effector from Pfm ICMP18803. The first column is an MLST tree using the core genes *gyrB*, *ropD*, *gapA*, *pgi*, *acnB*. The percentage homology at the nucleotide level is given for each result. Green boxes indicate a full-length coding sequence, and red boxes indicate the presence of a premature stop codon or a transposon insertion. Orange boxes indicate that it could not be ascertained whether the gene could produce a full-length protein, usually because the gene spanned two contigs in the short read assembly.

*P. syringae* pathovars can produce a range of toxic compounds, some of these, such as phaseolotoxin, coronatine and tabtoxin, have been well-characterised (Bender et al. 1999). Different biovars of Psa have been shown to produce various combinations of phaseolotoxin and coronatine (Fujikawa and Sawada 2019). Pfm does not produce any of the classic *P. syringae* phytotoxins, however analysis using AntiSMASH 6.0 (Blin et al. 2021) revealed nine secondary metabolite pathways (Table 2). The only characterised compound with a potential role in pathogenicity is mangotoxin, also known as Pseudomonas virulence factor (Carrión et al. 2012; Morgan et al. 2019). Other metabolites that might have a role in epiphytic fitness include various siderophores, the surfactant syringafactin and *N*-acetylglutaminylglutamine, which has a role in osmotic stress (Table 2). Of interest is the presence of the novel 3-thiaglutamate biosynthetic pathway, which uses a small peptide as a scaffold to synthesise a novel small molecule (Ting et al. 2019). This pathway has a limited distribution among *P. syringae* pathovars, with the majority of BLAST hits to *actinidifoliorum* and closely related *thea* pathovars.

The Pfm ICMP18803 genome was annotated using the NCBI genome workbench and resubmitted to GenBank (Kuznetsov and Bollin 2021). The availability of long reads greatly facilitated the correct annotation of effector genes and their relative location in the genome. Although Pfm is a poor pathogen of kiwifruit, a number of its reasonably extensive effector complement are present in Psa (McCann et al. 2013). This makes the comparison between the effector complements of Pfm and Psa a useful model for understanding how bacterial pathogens cause disease.

## Acknowledgments

We would like to thank Dr Nikki Freed and Annabel Whibley at Auckland Genomics (University of Auckland) for DNA sequencing using the MinION platform. We would also like to thank Drs Joanna Bowen and Erik Rikkerink (PFR) for reading this manuscript prior to publication.

## Author-Recommended Internet Resources

Oxford Nanopore Technologies: https://nanoporetech.com/

Geneious (Biomatters): https://www.geneious.com/

Flye: https://github.com/fenderglass/Flye

Pilon: https://github.com/broadinstitute/pilon

AntiSMASH: https://antismash.secondarymetabolites.org

